# Urbanization shapes the demographic history of a native rodent (the white-footed mouse, *Peromyscus leucopus*) in New York City

**DOI:** 10.1101/032979

**Authors:** Stephen E. Harris, Alexander T. Xue, Diego Alvarado-Serrano, Joel T. Boehm, Tyler Joseph, Michael J. Hickerson, Jason Munshi-South

## Abstract

How urbanization shapes population genomic diversity and evolution of urban wildlife is largely unexplored. We investigated the impact of urbanization on white-footed mice, *Peromyscus leucopus*, in the New York City metropolitan area using coalescent-based simulations to infer demographic history from the site frequency spectrum. We assigned individuals to evolutionary clusters and then inferred recent divergence times, population size changes, and migration using genome-wide SNPs genotyped in 23 populations sampled along an urban-to-rural gradient. Both prehistoric climatic events and recent urbanization impacted these populations. Our modeling indicates that post-glacial sea level rise led to isolation of mainland and Long Island populations. These models also indicate that several urban parks represent recently-isolated *P. leucopus* populations, and the estimated divergence times for these populations are consistent with the history of urbanization in New York City.

## INTRODUCTION

Urbanization is a particularly potent driver of environmental change around the world [1]. Understanding population genomic responses of organisms to human-driven change provides important context for predicting future evolutionary responses [2]. Using genome-wide SNP data, we investigate the effects of post-glacial environmental events and urbanization in the New York City (NYC) metropolitan area on historical demography of the white-footed mouse, *Peromyscus leucopus*. We examine the influence of climatic history over thousands of generations but also the effects of recent environmental events tens of generations in the past. This study is the first to examine the impact of urbanization on demographic history using patterns of genomic variation in wild populations.

NYC is particularly well suited for studies on urbanization because the city’s recent history of geological [3], ecological [4,5], and cultural [6,7] change has been meticulously recorded. NYC also has clearly defined urban green spaces that are delimited by anthropogenic and natural barriers, and occupied by independently-evolving populations of species with poor mobility through the urban matrix [8].

Natural barriers include the Hudson and East Rivers that separate the mainland portion of the city (i.e. Bronx) from Manhattan and Long Islands. The establishment of Long Island did not begin until the retreat of the late Wisconsin glacier that covered much of present-day NYC [9]. The glacier began retreating northward ~21,000 years before present (ybp) [10], and over the next few thousand years white-footed mice recolonized the region from southern refugia [11]. During this time, *P. leucopus* presumably maintained continuous populations until sea level rise separated Long Island from mainland NY between 12,000—15,000 ybp [10]. Except for occasional land-clearing by Native Americans, anthropogenic barriers were not erected until after European settlement of the area around 1600 CE [4]. During early phases of urbanization in NYC (1609-1790), green spaces within the city were parade grounds, cemeteries, farms, or private estates with highly manicured landscapes. In the mid-19^th^ century heavily used land plots, like present-day Prospect and Central Parks, were taken over by city officials and transformed for aesthetic purposes [12]. Private estates were also acquired by the NYC government and redesigned as vegetated parkland [13]. Remnant fauna in these parks, surrounded by a dense urban infrastructure, may have recovered from bottlenecks caused by urban fragmentation as the parks developed mature forests.

*P. leucopus* represents one of these remnant species, and we investigated the demographic history of populations occupying contemporary forest fragments in NYC and the surrounding area. *P. leucopus* are abundant across North America, have a typically short lifetime dispersal capability of ~100 m, prefer oak-hickory secondary forests, and consume a diet of arthropods, fruits, nuts, vegetation, and fungus. White-footed mice are abundant in small, fragmented urban forests [14–16], and exchange migrants only through vegetated corridors between isolated NYC parks [17]. Substantial genetic structure at microsatellite loci exists between NYC parks [8], and there is evidence of divergence and selection in genes underlying functional traits in urban populations [18].

In this study we estimated the demographic history of *P. leucopus* in NYC to test hypotheses about population expansion and divergence in response to urbanization. We used a genome-wide SNP dataset previously generated [19] from a double-digest restriction-site associated DNA sequencing (ddRADseq) [20] protocol. Loci came from 23 white-footed mouse populations (Fig 1) representative of a rural to urban gradient [19]. We used percent impervious surface cover and human population density around sampling sites as proxies for the extent of urbanization around each site (See Table 1 and Figure 1 in [19]). We then used sNMF version 0.5 [21] to examine population structure, and *TreeMix* [22] to build population trees and identify likely genetic clusters of *P. leucopus*. We used data from five populations of white-footed mice in NYC parks that showed evidence of genetic isolation and had relatively high urbanization metrics to test the hypothesis that temporal patterns of population isolation resulted from urbanization (Table 1, Figure S3). We estimated demographic parameters from the site-frequency-spectrum (SFS) using the composite-likelihood and coalescent simulation approach implemented in *fastsimcoal2 ver. 2.5.1* [23]. *Fastsimcoal2* efficiently calculates the approximate likelihood from unlinked SNP loci and accommodates complex demographic models. We used these estimates of effective population sizes, divergence times, migration, and population size changes to infer the influence of urbanization on the demography of these populations. Can we distinguish recent, human-driven demographic changes from older natural events under a complex model? See supplemental file 1 for full details on the methodology for this study.

**Figure 1.**
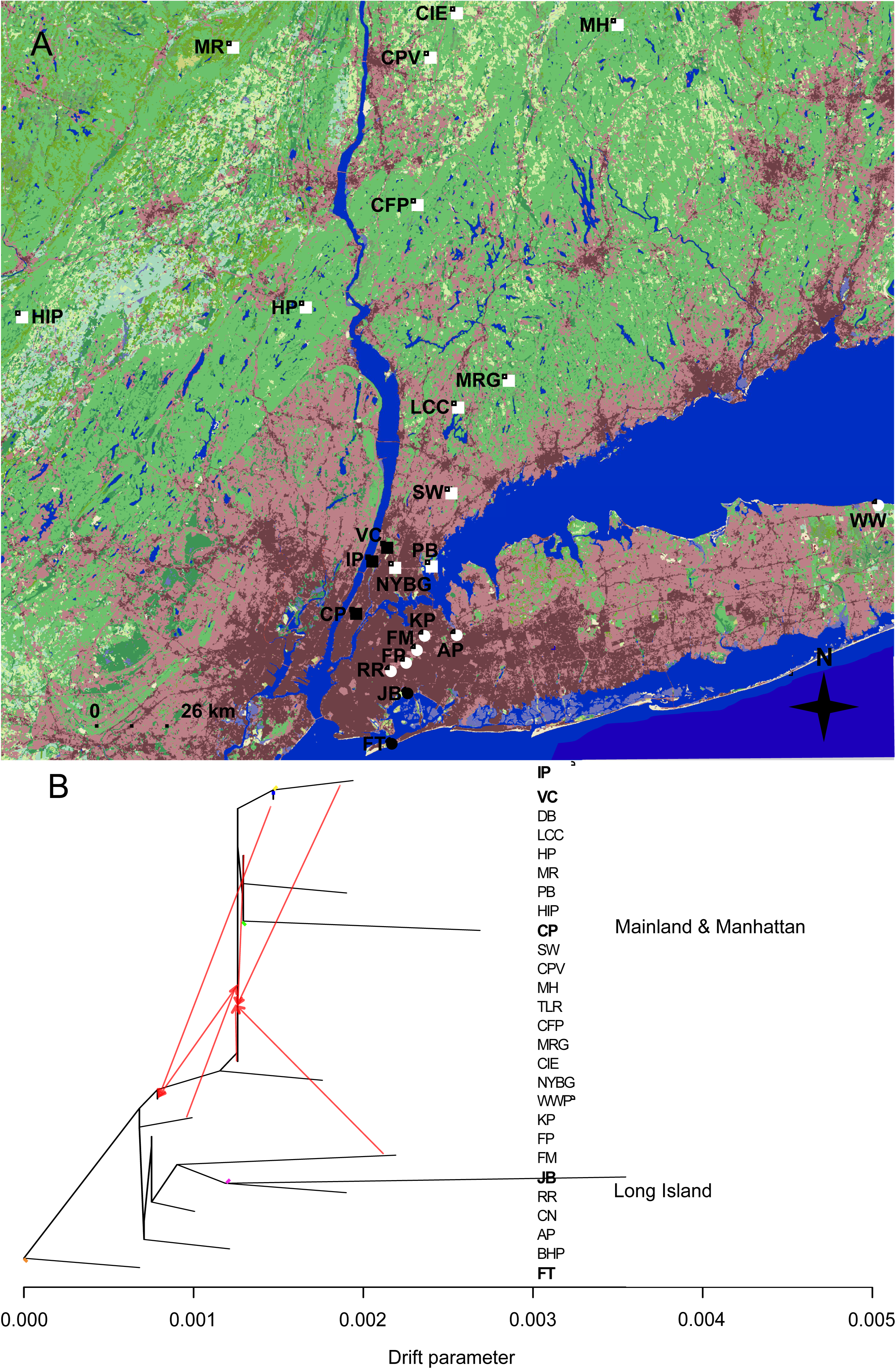
(A) Map of all sampling sites in NYC and the surrounding region. Colors correspond to the National Land Cover Database: Dark Red = Urban High Density Development; Light Red = Urban Medium to Low Density Development; Greens = Forested areas; Yellow = Grasslands. Squares = sampling sites from MM. Circles = sites from LI. Yellow shapes = sites used for urban population demographic analysis. (See Suppl File 1, Table S1 for full site names). (B) *TreeMix* population tree. Red arrows represent significant admixture using *TreeMix* and *f3* statistics. The drift parameter is plotted along the x-axis and represents the amount of genetic drift along the branch. Letters = sampling site codes (See Table S1 for full names, AP and CN were combined for all other analyses). Letters in bold and colored branches correspond to urban sampling sites described in Fig. 1A and show urban populations with relatively high levels of divergence to nonurban populations, as evidenced by long-branch lengths.

**Table 1.**
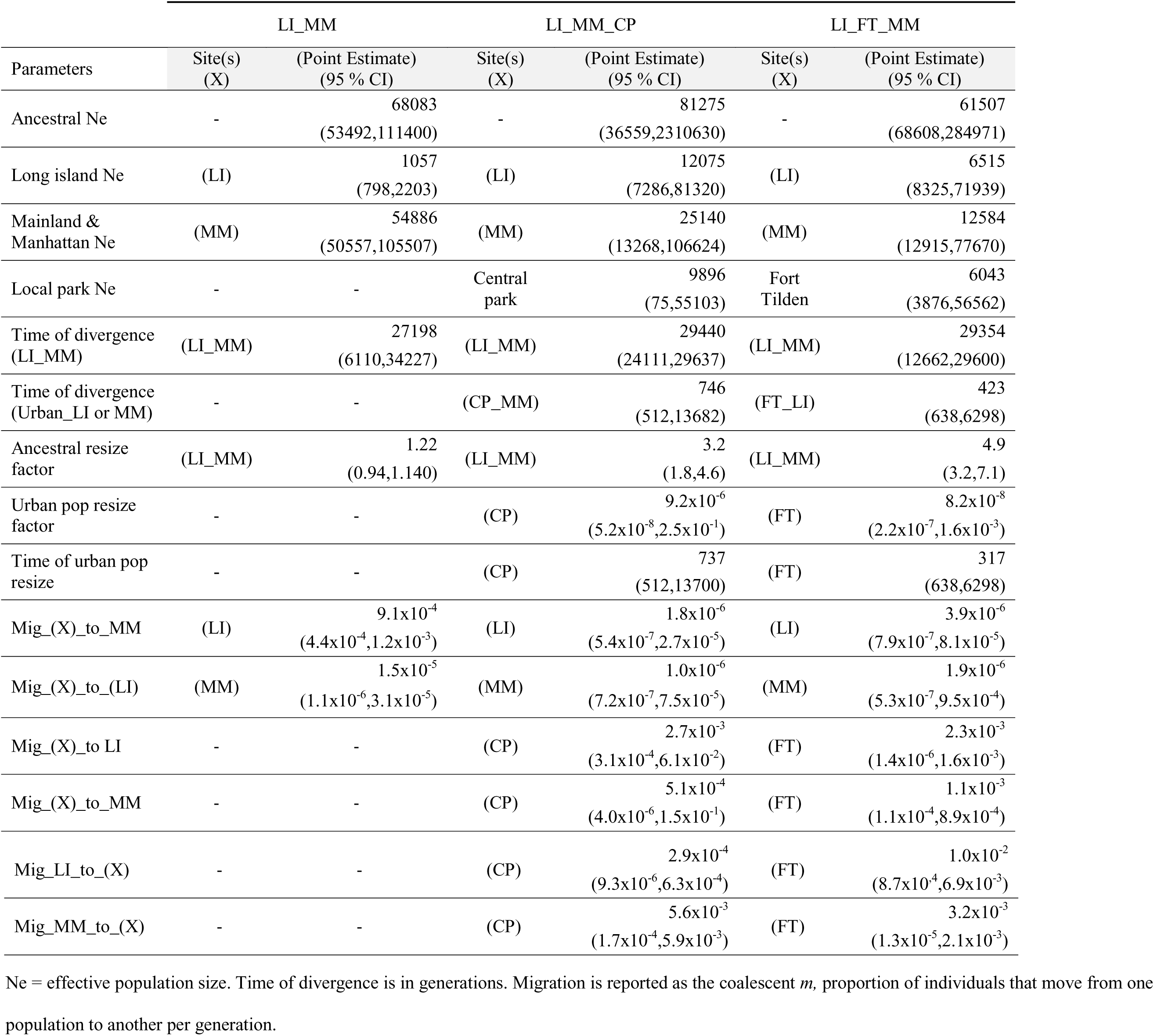
Inferred demographic parameters with 95% confidence values from parametric bootstrapping for the three main fastsimcoal2 model varieties. (See Table S2 for remaining models)

## RESULTS AND DISCUSSION

### Evidence for genetic structure and admixture

Our ddRAD dataset of 14,990 SNPs from 191 individuals sampled at 23 sites (mean of 8 ± 0.17 individuals / site) [19] captured sufficient genetic variation to estimate the post-glacial demographic history of white-footed mouse populations in the NYC metropolitan area. Before inferring demography, a sparse non-negative matrix factorization approach (*sNMF*, Frichot *et al*. 2014) supported assignment of individuals into two main groups separated by the East River and Long Island Sound: 1) Mainland & Manhattan (MM) and 2) Long Island (LI; Fig. S1). Population trees from *TreeMix* [22] supported the patterns inferred using sNMF. *TreeMix* also indicated that several urban parks contain recently-fragmented populations (Fig 1B) with no evidence of admixture with other sites (Supplemental File 2). When assigning individuals to populations for demographic model development, we compared our results to those of a previous study that examined population structure using genome-wide loci [19]. Genetically differentiated populations included Central (area: 344.05 ha, 2 km buffer % impervious surface & human population size: 60.2, 351698.8), Inwood (79.21 ha, 2 km buffer % impervious surface & human population size: 30, 121354.2), and Van Cortlandt (433.15 ha, 2 km buffer % impervious surface & human population size: 27.7, 77541.7) Parks in MM (790,142 ha); and Jamaica Bay (263.38 ha, 2 km buffer % impervious surface & human population size: 3.2, 1438.4) and Fort Tilden (248.96 ha, 2 km buffer % impervious surface & human population size: 8.5, 2357.5) in LI (362,900 ha). These urban parks are all large, extensively vegetated, and surrounded by dense urban development (Fig. 1A). No rural sampling locations exhibited patterns consistent with genetically isolated populations, suggesting the parks above were isolated due to urbanization.

### *P. leucopus* population history during recent urbanization in NYC

Inferred parameter estimates exhibit a consistent signal of an older split between LI and MM populations in line with geologic records followed by recent divergence of NYC park populations (Figure S2). Models had tight confidence intervals around divergence times for MM and LI (~13,600 ybp, Fig S2-E) except for the two-population model. The two population model had the lowest likelihood and this result may reflect the relatively poor fit of the model. Divergence was followed by a strong population contraction (Table 1, Fig. S3). These divergence estimates concur with geologic records that date the separation of Long Island and the Mainland from ~13,000 – 15,000 ybp [24].

Our other demographic models examined whether contemporary urban populations diverged from MM or LI within the historical timeframe of urbanization in NYC. In 1609, shortly after European arrival, only 1% of the Manhattan landscape was urbanized. Over the next 400 years, humans converted 97% of natural green spaces to human use [4]. Urban populations experienced strong population bottlenecks at the time of divergence (except Jamaica Bay) and the inferred time of divergence was always within the 400-year window of European settlement (Table 1). While 400 years, representative of ~800 *P. leucopus* generations assuming a generation time of 0.5 years, is relatively recent, detailed demographic inference over very recent time scales is possible with adequately large genomic datasets [23]. Additionally, many point estimates for urban park divergence are in line with the founding of urban parks in NYC (282 ybp – present, Table 1). These results indicate that isolation in urban fragments was sufficiently strong to impact the evolutionary history of urban fauna.

We detected bottlenecks immediately after isolation of urban populations, suggesting that a small remnant population within these parks at the time of the bottleneck provided most of the urban genetic variation found today. Our inferred migration rates between all populations were high and variable, but we estimated consistent patterns of low migration between MM and LI, and asymmetrical migration of individual mice from MM into urban populations (Table 1). Despite asymmetrical gene flow, urban parks consistently showed a signal of some emigration to LI or MM, suggesting that urban parks contain stable, though relatively small populations. However, given the extremely recent divergence times, these high migration rates could be due to retained ancestral polymorphisms from incomplete lineage sorting or geographic structure that are difficult to distinguish from admixture [25]. It is important to note that allelic dropout in ddRADSeq data from mutations in cut sites can affect demographic analyses, but using a minimum coverage cutoff and restricting the amount of missing data can mitigate these effects (Supplementary File 1).

## CONCLUSIONS

Our results show that geography, geologic events, and human-driven habitat change have left a detectable genomic signature in NYC’s white-footed mouse populations. Patterns of genetic variation and population structure reflect past demographic processes [26], and genome-wide SNPs generated from ddRADseq provided enough information to distinguish recent demographic events from past geological processes. Our demographic models estimated divergence times and migration patterns that are consistent with the known geologic and historical record of NYC. This study is the first to use population genomic modeling to estimate the demographic impact of urbanization on wild populations.

## DATA ACCESSIBILITY

Illumina sequencing reads from Munshi-South *et al*. (2016) have been deposited in NCBI's Short-read Archive (SRA) under accession number SRP067131. The VCF file of SNP genotypes used here and in Munshi-South *et al*. (2016) is available on the Dryad digital repository at http://dx.doi.org/10.5061/dryad.d48f9

## ETHICS STATEMENT

All animal handling procedures were approved by the Institutional Animal Care and Use Committee (IACUC) at Fordham University (Protocol No. JMS-13-03). Samples were collected with permission from the New York State Department of Environmental Conservation, New York City Department of Parks and Recreation, New York Botanical Garden, and the Connecticut Department of Energy and Environmental Protection.

## ACKNOWLEDGMENTS

We thank the Hickerson lab for access to space and productive conservations on this topic, and Laurent Excoffier for guidance on the demographic inference. The Handling Editor and three anonymous reviewers for *Biology Letters* provided many helpful suggestions for improving the manuscript.

## AUTHOR CONTRIBUTIONS

S.E. Harris conceived and designed the study and conducted analyses and interpretation of the data. A.T. Xue, D. Alvado-Serrano, J.T. Boehm, T. Joseph, and M. J. Hickerson conducted analyses and interpretation of the data. J.Munshi-South conceived and designed the study, acquired the samples and genetic data, and conducted analyses and interpretation of the data. All authors drafted the article and revised it critically for important intellectual content. All authors approved the final version of the published manuscript, and agree to be held accountable for all aspects of the work herein.

## METHODS

### Sampling and DNA extractioxn

During two previous studies [1,2] we sampled individual white-footed mice between 2009 and 2013 from 23 separate localities that were used to generate the genomic data used in this study. Sites were chosen to represent a rural to urban gradient (Fig. 1). Rural sites were defined as large tracts of relatively undisturbed natural habitat, and urban sites were fragmented habitat surrounded by urban infrastructure and impervious surface. Urbanization was also quantified by determining the percent impervious surface and human population size in a 2 km buffer surrounding each park (See Table 1 and Figure 1 in [2]). For all sampling locations, we trapped individuals over a period of 1-3 nights each. At each site, we set between one and four 7x7 m transects of Sherman live traps (7.62 cm x 7.62 cm x 22.86 cm), depending on the total area of each sampling site. We weighed, sexed, and took morphological (ear length, tail length, hind foot length, total body length) measurements for all individual mice. At all sites except Central Park, Flushing Meadow, New York Botanical Garden, Brook Haven Park & Wild Wood Park, High Point Park, and Clarence Fahnestock Park, we collected tissue by taking 1 cm tail clips, placing in 80% ethanol, and storing at −20° C in the laboratory. Individual mice were then released. For these other six sites, we used previously-collected liver samples stored in RNAlater (Ambion Inc., Austin, TX) at −80° C. We extracted genomic DNA using standard extraction protocols, quantified the yield, and checked quality before genomic sequencing library preparation. See methods in (Munshi-South *et al*. 2016) for full details. All animal handling procedures were approved by the Institutional Animal Care and Use Committee at Brooklyn College, CUNY (Protocol Nos. 247 and 266) and by Fordham University’s Institutional Animal Care and Use Committee (Protocol No. JMS-13-03).

### RAD sequencing and SNP calling

We initially sequenced 233 individual white-footed mice but retained 191 *P. leucopus* individuals from 23 sampling sites for the genome-wide SNP dataset after filtering out close relatives and low-quality samples [2]. Briefly, we followed standard protocols for ddRADseq presented in Peterson *et al*. (2012), starting with DNA extraction using Qiagen DNEasy kits with an RNAse treatment. Next we used a combination of the enzymes, SphI-HF and MluCI to generate similarly sized DNA fragments. Using two restriction enzymes increases the probability of generating the same RAD loci across all samples. We specifically chose SphI-HF and MluCI to generate ~50,000 RAD loci; this estimate was based on the number of fragments produced using the same REs on a related species (*Mus musculus*). We cleaned the digested DNA with Ampure XP beads and then ligated unique barcodes to each individual sample. We then pooled samples in groups of 48 and used a Pippin Prep for precise DNA fragment size excision from gels and Phusion high-fidelity PCR to amplify fragments and add Illumina indexes and sequencing primers. The resulting fragments were sent to the NYU Center for Genomics and Systems Biology for 2x100 bp paired-end sequencing in three lanes of an Illumina HiSeq 2000. We checked initial quality of the raw reads using FastQC. Subsequent primer removal, low-quality nucleotide trimming, and *de novo* SNP calling was conducted using the Stacks 1.21 pipeline [4]. We called and filtered SNPs in Stacks using default setting except for requiring that loci occur in 22 / 23 sampling sites, and within each site, occur in at least 50% of individuals. We chose a random SNP from each RAD tag to avoid linkage between loci. Additionally, we removed individuals if they had too few reads resulting in extremely small SNP datasets or if they showed high levels of relatedness to other white-footed mice sampled. These filters resulted in 14,990 SNPs in the final dataset that we used for demographic modeling.

### Population structure and migration

We investigated observed patterns of genetic diversity to define evolutionary clusters that could be used to inform demographic modeling of *P. leucopus* populations in the NYC region. We examined population structure and evidence of migration among all 23 sampling sites. The program *TreeMix* [5] was used to build population trees and identify migration events. *TreeMix* infers populations splitting and mixing using allele frequencies from large genomic datasets. Using a composite likelihood approach given allele frequency data, TreeMix returns the most likely population tree and admixture events given a user-specified number of admixture events. The number of admixture events tested ranged from 0 - 12 while the rest of the parameters used default settings. P-values were generated for each admixture event and comparisons made between all trees. We confirmed admixture between populations by running *f3* three-population analyses in Treemix. These statistics assess admixture between populations by identifying correlations between allele frequencies that do not fit the evolutionary history for that group of three populations. We used 500 bootstrap replicates to assess significance of *f3* statistics.

We also used sNMF version 0.5 [6] to examine population structure sNMF explores patterns of genetic structure by assigning individual ancestry coefficients using sparse nonnegative matrix factorization. sNMF does not make any model assumptions like requiring populations to be in Hardy-Weinberg and linkage equilibrium [6], as opposed to other likelihood models like STRUCTURE [7]. For the number of putative ancestral populations tested, we chose a range from K = 1 to K = 11 using default parameters, with 10 replicate runs for each value of K. We chose 11 as an initial maximum because there were at least nine urban parks without any vegetated corridors between them (parks in Queens, NY have a small greenway connecting them) plus rural LI and rural Mainland. There was not a pattern supporting a higher number of clusters so we did not analyze K > 11. We ran sNMF on the full 14,990 SNP dataset (≤ 50% of SNPs missing per population) and on a more conservative dataset with only ≤ 15% of SNPs missing per population. sNMF imputes missing genotypes by resampling from the empirical frequency at each locus [6], and using fewer missing data ensured any inferred population structure was not due to incorrectly imputed genotypes (Fig. S1). To infer the most likely number of ancestral populations, each model run generates a cross-entropy estimation based on ancestry assignment error when using masked genotypes. The model with the smallest cross-entropy score implies it is the best prediction of the true number of K ancestral populations [6].

### Demographic inference from genome-wide site frequency spectra

To reduce model complexity for demographic inference, we grouped individuals into the minimum number of populations representing unique evolutionary clusters. Global analyses in TreeMix and sNMF showed the highest support for two populations split by the East River, and hierarchical analyses using discriminate analysis of principal components [2] showed support for isolated urban populations. Collectively, results suggested a minimum of seven putative populations captured most of the genetic variation between populations (Mainland & Manhattan: MM, Long Island: LI, Central Park: CP, Van Cortlandt Park: VC, Inwood Hill Park: IP, Jamaica Bay: JB, Fort Tilden: FT, Fig. 1). Along with hierarchical population structure results, we chose several of the urban populations to include in demographic modeling based on the size of the park, the relative isolation of the park due to urbanization, and the population density of white-footed mice in the park. We generated the multi-population site frequency spectrum (MSFS) for subsets of populations to test specific demographic history scenarios. We used the *dadi.Spectrum.from_data_dict* command implemented in *dadi* [8] to generate the MSFS. When we created the SNP dataset, we required a SNP to occur in ≥ 50% of individuals from each population, so the MSFS was down-projected to 50% to ensure the same number of individuals for all loci [8]. Once the MSFSs were generated, we used the software program fastsimcoal2 ver 2.5.1 [9] for demographic inference. *Fastsimcoal2* (fsc2) uses a composite multinomial likelihood approach to infer demographic histories from the site frequency spectrum generated from genomic scale SNP datasets. The expected SFS under user defined demographic scenarios is obtained using coalescent simulations. *Fastsimcoal2 ver. 2.5.1* contained a bug that miscalculated Max Observed Likelihood values if the SFS contained non-integers, leading to Maximum Estimated Likelihoods that are higher than Max Observed Likelihoods. Our data did not display this symptom and when point estimates were re-run using the latest version (fsc25), there was no impact on the values used here and on the final conclusions presented in the main text.

We tested demographic histories under a scenario of population isolation with migration (IM). We compared inferred parameters between six hierarchical IM models (Fig. S3) using the same dataset. There was one two-population IM model (seven free parameters) to test older divergence patterns between MM and LI suggested from the geologic record. The remaining five models were three-population IM models (15 free parameters each) testing for recent urban population divergence. We chose to run separate models investigating each urban population separately in order to avoid inconsistencies from over-parameterization. For these remaining models we considered an ancestral population that split at time *T*_*div1*_ and then an urban population that split more recently at time *T*_*div2*_. For *T*_*div1*_ we included a range of divergence times based on the LGM of the Wisconsin glacier, ~18,000 ybp. For *T*_*div2*_ we considered divergence times incorporating the timeframe of urbanization in NYC, ~400 ybp. We allowed for migration between all populations, and tested occurrences of population bottlenecks when urban isolation was incorporated into the model. During likelihood calculation, a conditional maximization algorithm (ECM) is used to maximize the likelihood of each parameter while keeping the others stabilized. This ECM procedure runs through 40 cycles where each composite-likelihood was calculated using 100,000 coalescent simulations. While increasing the number of simulations can increase precision, accuracy does not significantly increase past 100,000 simulations [9]. Additionally, in order to avoid likelihood estimates that oversample parameter values at local maxima across the composite likelihood surface, we ran 50 replicates with each starting from different initial conditions. We chose the replicate with the highest estimated maximum likelihood score for each model. Using parametric bootstrapping, we generated confidence intervals for the most likely inferred demographic parameters generated. The SFS was simulated with the parameter values from the highest likelihood model and then new parameter values reestimated from the simulated SFS. We ran 100 parametric bootstraps. To find consistent signals of divergence that could be attributed to urbanization, we compared parameter values and overlapping confidence intervals between models.

## DEMOGRAPHIC INFERENCE

Parameters were allowed to vary in demographic modeling using fastsimcoal2, but all six models converged on similar parameter values estimated from the observed MSFS. Parameter estimates with the highest likelihood generally fell within the upper and lower bounds generated from parametric bootstrapping (Fig. S2, Table 1). The first two-population model tested divergence time, effective population size, and migration rates between MM and LI populations (LI_MM, Fig. S3). The divergence time for the MM and LI split was inferred to be 13,599 ybp and the effective density of white-footed mice in MM (*N*_*E*_ / size, 0.069 mice/ha) 23x larger than LI (*N*_*E*_ / size, 0.003 mice/ha) (Table 1). Divergence times are based on a generation time of 0.5 years for *Peromyscus leucopus*. Migration was also inferred to be low (< 1 individual per generation) between MM and LI (Table 1).

The inferred demography for the more complex three-population models generally supported results from the two-population model. The first two complex models both estimated the divergence between MM and LI, but one model tested for divergence of JB and LI after the MM and LI split (LI_JB_MM, Fig. S3) while the other model tested divergence between FT and LI after the MM and LI split (LI_FT_MM, Fig. S3). This model also tested the likelihood of a bottleneck event when FT and JB, both urban populations, diverged. We set up the other three complex models in an identical fashion, except we tested the urban populations of CP (LI_MM_CP, Fig. S3), VC (LI_MM_VC, Fig. S3), or IP (LI_MM_IP, Fig. S3) for divergence from MM after the MM and LI split. Point estimates for demographic parameters converged on similar values and generally fell within the 95% confidence limits from parametric bootstrapping (Fig. S2, Table 1). The average divergence time for MM and LI was 14,679 ybp, SD = 956.19. Similar to the two-population model, the MM *N_E_* was larger than the LI *N*_*E*_. While the effective population sizes for the urban parks were smaller than *N*_*E*_ for MM or LI, the urban populations contained much higher effective population densities (CP: 28.7 mice / ha, IP: 79.21 mice / ha, JB: 23.8 mice / ha, FT: 24.3 mice / ha, VC: 0.36 mice / ha) than either MM (0.069) or LI (0.003). The individual urban populations all had *N*_*E*_ values 10x smaller than MM, but often similar to LI. The divergence time for the five tested urban populations, even with variation in number of generations per year, was consistent with the timeframe of urbanization (mean divergence = 233 ybp; SD = 164.5). Our demographic models proved to be rather robust in returning reasonable parameter values with consistent convergence to similar values across replicates. Although wide confidence intervals on many parameters suggest low resolution in inferring parameter values given the model and data, they are likely a consequence of the complexity of the model given the number of parameters and wide parameter ranges. The narrow confidence intervals on other parameters suggest that these inferences reliably capture important aspects of the true demographic history of white-footed mice in NYC, especially given the often biologically unrealistic parameter search space (See est files). One limitation in using Radseq data for demographic analysis is the effect that allelic dropout has on genetic variation. Mutations can accumulate in RE cut sites causing the loss of loci with potentially higher mutation rates [10]. Thus, ddRADSeq may underestimate genetic diversity [11] and inflate sequence divergence [12,13]. We addressed this limitation by setting a minimum coverage limit and minimizing missing data that might be caused by allelic dropout [11]. We also used multiple other methods including discriminant analysis of principle components, sparse non-negative matrix factorization, classical population genetics statistics, and composite likelihood for population tree inference, to confirm our demographic analysis findings and inform our demographic modeling.

